# Wave-like activity patterns in the neuropil of striatal cholinergic interneurons in freely moving mice represent their collective spiking dynamics

**DOI:** 10.1101/404467

**Authors:** Rotem Rehani, Yara Atamna, Lior Tiroshi, Wei-Hua Chiu, José de Jesús Aceves Buendía, Gabriela J. Martins, Gilad A. Jacobson, Joshua A. Goldberg

## Abstract

Cholinergic interneurons (CINs) are believed to form synchronous cell assemblies that modulate the striatal microcircuitry and possibly orchestrate local dopamine release. We expressed GCaMP6s, a genetically encoded calcium indicator (GECIs), selectively in CINs, and used microendoscopes to visualize the putative CIN assemblies in the dorsal striatum of freely moving mice. The GECI fluorescence signal from the dorsal striatum was composed of signals from individual CIN somata that were engulfed by a widespread fluorescent neuropil. Bouts of synchronous activation of the cholinergic neuropil revealed traveling-wave-like patterns of activity that preceded the signal from individual somata. To investigate the nature of the neuropil signal and why it precedes the somatic signal, we target-patched GECI-expressing CINs in acute striatal slices in conjunction with multiphoton imaging or wide-field imaging that emulates the microendoscopes’ specifications. The ability to detect fluorescent transients associated with individual action potential was constrained by the long decay constant of GECIs (relative to common inorganic dyes) to slowly firing (< 2 spikes/s) CINs. The microendoscopes’ resolving power and sampling rate further diminished this ability. Additionally, we found that only back-propagating action potentials but not synchronous optogenetic activation of thalamic inputs elicited observable calcium transients in CIN dendrites. Our data suggest that only bursts of CIN activity (but not their tonic firing) are visible using endoscopic imaging, and that the spatiotemporal neuropil patterns are a physiological measure of the collective recurrent CIN network spiking activity.

**Significance Statement:** Cholinergic interneurons (CINs) are key modulators of the striatal microcircuitry that are necessary for assigning action value and behavioral flexibility. We present a first endoscopic imaging study of multiple molecularly identified CINs in freely moving mice. We reveal the presence of traveling-wave-like activity in the CIN neuropil. We then use *ex vivo* electrophysiological and imaging techniques to show that the neuropil signal is the integrated fluorescence arising from the axo-dendritic arbors of CINs dispersed throughout the striatum. We conclude that the neuropil signal acts as a mean-field readout of the striatal CIN network activity.

## Introduction

Striatal cholinergic interneurons (CINs) are the main population of tonically active neurons (TANs) whose pause response is associated with the presentation of reward or with stimuli that are associated with reward (Anderson, 1978; Kimura et al., 1984; Aosaki et al., 1994; Raz et al., 1996). It was hypothesized long ago that CINs form synchronous striatal cell assemblies during the pause responses (Graybiel et al., 1994). These assemblies collectively modulate neuronal excitability, synaptic transmission and synaptic plasticity in the striatal microcircuitry (Calabresi et al., 2000; Pakhotin and Bracci, 2007; Pisani et al., 2007; Goldberg et al., 2012; Plotkin and Goldberg, 2018). This hypothesis has been buttressed by recent *ex vivo* data showing that synchronous activation of CINs can lead to the release of dopamine, GABA and glutamate in the striatum (Cachope et al., 2012; English et al., 2012; Threlfell et al., 2012; Chuhma et al., 2014; Nelson et al., 2014). Multiple electrode recordings in primates have shown that TANs exhibit some degree of synchrony which increases in experimental parkinsonism (Raz et al., 1996; Apicella et al., 1997; Goldberg et al., 2002; Ravel et al., 2003; Goldberg et al., 2004). With the advent of genetically encoded calcium indicators (GECIs) it is now possible to conduct longitudinal studies on large assemblies of molecularly identified neurons, such as the CINs, in freely moving mice performing self-initiated movements or undergoing training (Mank and Griesbeck, 2008; Ghosh et al., 2011; Lin and Schnitzer, 2016). In most GECI experiments, the images are composed of fluorescence that arises from the somata of the targeted neurons and due to background fluorescence.

In most studies the background activity is presumed to arise from out-of-focus neurons, hemodynamics (due to the auto-fluorescence of blood vessels) and other artifacts such as motion or photo-bleaching of dyes (Pnevmatikakis et al., 2016; Zhou et al., 2018). In the case of microendoscopic imaging with GRIN lenses implanted deep in the brain and where diffuse light contamination can be minimized, the contribution of hemodynamics is likely to be less of an issue (Ma et al., 2016). Nevertheless, most recent studies from various groups that conduct microendoscopic imaging have adopted a strategy that calls for the removal of the background signal in order to extract a cleaner neuronal signal. Two approaches have been used to subtract the background signal. The first is a heuristic and involves estimating the background fluctuations in the vicinity of a given soma. This signal (or a weighted version of it) is subtracted from the somatic signal (Klaus and Plenz, 2016; Stamatakis et al., 2018; Zhou et al., 2018). The logic is simple. Due to the substantial depth-of-field of the imaging system, any signal observed in the vicinity is likely to contaminate the pixels in the soma and must therefore be subtracted. The other approach builds on the fact that the background signal tends to be highly correlated in space, and therefore lends itself to being modeled as a global background signal composed of a DC term plus some low spatial frequency components. By adding this assumption to an assumption regarding the parametric exponential shape of calcium events that accompany individual spikes, this approach has yielded sophisticated algorithms which simultaneously estimate independent neuronal sources while subtracting a global model of the background (Klaus and Plenz, 2016; Stamatakis et al., 2018; Zhou et al., 2018).

However, when transfecting the neurons, the fluorescent proteins are expressed in all compartments of the neurons including the axon and dendrite (Kerr et al., 2005; Lee et al., 2017). When considering the known anatomy of CINs that possess very intricate axonal and dendritic arbors (Chang et al., 1982; DiFiglia, 1987; Wilson et al., 1990; Kawaguchi, 1992), it stands to reason that a large component of the background signals is not out-of-focus somatic signals but rather calcium influx due to propagation and/or back-propagation action potentials throughout the CINs’ axonal and dendritic arbors, respectively. Under these circumstances, these background signals should be tightly related to the network state of the CIN network, implying that the background signal could provide a physiological readout of the entire CIN network.

In the current study, we report – for the first time – the presence of traveling wave-like activity in the GECI signals from striatal cholinergic neuropil in freely moving mice imaged with microendoscopes. In order to better understand the origin of the neuropil signal we use wide field (“one-photon”) and two-photon laser scanning microscopy (2PLSM) imaging of GECI signals from CINs in acute striatal slices. The combination of approaches leads us to the conclusion that to a large degree the spatiotemporal patterns observed in the cholinergic neuropil arise from back-propagating APs (bAPs) in the dendritic arbors. As a sum of many cholinergic processes the background activity, like local field potentials (LFPs), represents a readout of the collective discharge of CINs.

## Materials and Methods

### Animals

Experimental procedures adhered to and received prior written approval from the The Hebrew University’s Institutional Animal Care and Use Committee. Two-to-nine-months-old choline acetyltransferase-cre dependent (ChAT-IRES-Cre) transgenic mice (stock number 006410; Jackson Laboratories, Bar Harbor, ME, USA) of both sexes were used in the experiments.

### Stereotaxic injection of adeno-associated viruses and ChR2

Mice were deeply anesthetized with isoflurane in a non-rebreathing system (2.5% induction, 1–1.5% maintenance) and placed in a stereotaxic frame (model 940, Kopf Instruments, Tujunga, CA, USA). Temperature was maintained at 35°C with a heating pad, artificial tears were applied to prevent corneal drying, and animals were hydrated with a bolus of injectable saline (5 ml/kg) mixed with an analgesic (5 mg/kg carpofen). To calibrate specific injection coordinates, the distance between bregma and lambda bones was measured and stereotaxic placement of the mice was readjusted to achieve maximum accuracy. A small hole was bored into the skull with a micro drill bit and a glass pipette was slowly inserted at the relevant coordinates in aseptic conditions. To minimize backflow, solution was slowly injected and the pipette was left in place for 7 min before slowly retracting it.

A total amount of 400 nl of an adeno-associated virus serotype 9 harboring GCaMP6s construct (AAV9-syn-flex-GCaMP6s; > 2.5 × 10^13^ viral genome/ml; University of Pennsylvania Vector Core, catalog number AV-9-PV2824) was injected into the dorsal striatum under aseptic conditions. The coordinates of the injection were as follows: anteroposterior, +0.5 mm; mediolateral, +2.3 mm; and dorsoventral, −2.8 mm, relative to bregma using a flat skull position (Paxinos and Franklin, 2012).

For thalamic expression of ChR2 a total of 250 nl of an adeno-associated virus serotype 9 harboring ChR2 construct (AAV9-hSyn-ChR2-eGFP; > 2.5 × 10^13^ viral genome/ml; University of Pennsylvania Vector Core, catalog number AV-9-26973P) was injected into the caudal intralaminar nucleus (ILN) of the thalamus to transfect the parafascicular nucleus (PfN) neurons under aseptic conditions. The coordinates of the injection were as follows: anteroposterior, −2.3 mm; mediolateral, +0.65 mm; and dorsoventral, −3.35 mm, relative to bregma using a flat skull position (Paxinos and Franklin, 2012; Ellender et al., 2013). Two to three weeks after viral injection mice were used for experiments.

### Gradient refractive index (GRIN) lens implantation

One week after the stereotaxic injection, mice were deeply anesthetized with isoflurane in a non-rebreathing system and placed in the stereotaxic frame, as described above (in some cases, a bolus of ketamine (32 mg/kg)-xylasine (3.2 mg/kg) was injected initially to stabilize the preparation for and induction of anesthesia). A hole slightly wider than the 1mm diameter (4 mm long) GRIN lens was drilled into the skull in aseptic conditions. We aspirated cortex with a 27-30 G needle to a depth of approximately 2 mm (just past the corpus callosum) and then fit the lens in snugly and (dental-) cemented it into place together with a head bar for restraining the mouse when necessary. One week later we attached a baseplate to guarantee that the endoscope will be properly aligned with the implanted GRIN lens. To ensure light impermeability, the dental cement was mixed with coal powder and painted with black nail polish. Two weeks later, the freely-moving mice were imaged in a behavior chamber lit by diffuse infrared light.

### Microendoscopic Imaging

Microendoscopes (*nVista*, Inscopix, Palo Alto, CA, USA) sampled an area of approximately 600 µm by 900 µm (pixel dimension: 1.2 µm) at 10 frames/sec. Movies were motion corrected with the Inscopix Data Processing Software (IDPS) suite. Motion-corrected movies and electrophysiological data were analyzed, and curve fitting was performed using custom-made code in MATLAB (MathWorks, Natick, MA, USA). We extracted fluorescence changes over time (*∆F/F*) such that 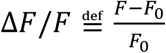. *F* is the raw fluorescent values recorded; *F*_0_ denotes the minimal averaged fluorescence in 1.5 sec-overlapping 3 sec periods throughout the measurement. Mice with weak transfection or too few somata were discarded. Somata were identified from a long-term temporal maximum projection of the *∆F/F* signal, and Regions-of-Interest (ROIs) were marked manually to engulf the somatic area. The annulus of each ROI was defined as a ring of pixels with the same area as the ROI, whose inner diameter is the distance of the point on the border of the ROI that is farthest away from the center-of-mass of the ROI plus 5 additional pixels. These annuli were also used to estimate the collective neuropil signal. We used an alternative scheme to estimate the collective signal. We considered all parts of the image that were devoid of somata and their surrounding 40 pixels. We then chose 100 circular ROIs with a radius of 5 pixels randomly located within this region.

To determine the temporal relationship between the somatic and annular signals, we detected major events in the somatic signal and averaged both the somatic and annular signals around these times. Signals were first averaged over all of the events in each soma-annulus pair, and the resulting traces were then averaged over the pairs. Our criterion for choosing the pairs was that they must display at least 5 events that did not contain another event, either in the soma or the annulus, within the 3 seconds following the peak.

### Slice preparation

Two to three weeks following AAV injections, mice were deeply anesthetized with intraperitoneal injections of ketamine (200 mg/kg) – xylazine (23.32 mg/kg) and perfused transcardially with ice-cold modified artificial CSF (ACSF) oxygenated with 95% O_2_–5% CO_2_ and containing the following (in mM): 2.5 KCl, 26 NaHCO_3_, 1.25 Na_2_HPO_4_, 0.5 CaCl_2_, 10 MgSO_4_, 10 glucose, 0.4 Ascorbic acid, and 210 sucrose. The brain was removed and blocked in the sagittal plane and sectioned at a thickness of 240 μm in ice-cold modified ACSF. Slices were then submerged in ACSF, bubbled with 95% O_2_-5% CO_2_, and containing the following (in mM): 2.5 KCl, 126 NaCl, 26 NaHCO_3_, 1.25 Na_2_HPO_4_,2 CaCl_2_, 2 MgSO_4_, and 10 glucose and stored at room temperature for at least 1 h before recording and/or imaging.

### Slice visualization and data collection – wide-field imaging

Slices were transferred to a recording chamber mounted on an Olympus BX51 upright, fixed-stage microscope and perfused with oxygenated ACSF at room temperature. A 20X, 0.5 NA water immersion objective was used to examine the slice using Dodt contrast video microscopy.

#### Electrophysiology

Electrophysiological recordings were obtained with a Multiclamp 700B amplifier (Molecular Devices, Sunnyvale, CA). The junction potential, which was 7–8 mV, was not corrected. Signals were digitized at 10 kHz. Patch pipette resistance was typically 3– 4 MΩ when filled with the recording solution. For calcium imaging experiments in conjunction with current-clamp recordings the pipette contained the following (in mM): 130 K-gluconate, 6 KCl, 8 NaCl, 10 HEPES, 2Mg_1.5_ATP, pH 7.3 with KOH (280–290 mOsm/kg).

#### One-photon wide-field calcium imaging

Optical measurements were made using 470 nm LED illumination (Mightex, Toronto, ON, Canada) and a cooled EM-CCD (Evolve 512 Delta, Photometrics, Tucson, AZ, USA). Frames were 512×512 pixels, pixel size was 0.4 µm with no binning and frame rate was 5-10 Hz. Regions of interest were marked manually offline and fluorescent traces were extracted.

Optical and electrophysiological data were obtained using the custom-made shareware package WinFluor (John Dempster, Strathclyde University, Glasgow, Scotland, UK), which automates and synchronizes the imaging signals and electrophysiological protocols. *∆F/F* (for all acute slice experiments) was calculated as defined above with *F*_0_ defined as the minimal value attained during the trace.

### Slice visualization and data collection – 2PLSM

The slices were transferred to the recording chamber of FemtO2D-Galvo scanner multiphoton system (Femtonics Ltd., Budapest, Hungary) and perfused with oxygenated ACSF at room temperature. A 16X, 0.8 NA water immersion objective was used to examine the slice using oblique illumination.

#### Electrophysiology

Electrophysiological recordings were obtained with a Multiclamp 700B amplifier (Molecular Devices, Sunnyvale, CA). The junction potential, which was 7–8 mV, was not corrected. Signals were digitized at 40 kHz. Patch pipette resistance and solution were as described for one-photon experiments.

#### 2PLSM calcium imaging

The 2PLSM excitation source was a Chameleon Vision 2 tunable pulsed laser system (680–1,080 nm; Coherent Laser Group, Santa Clara, CA). Optical imaging of GCaMP6s signals was performed by using a 920-nm excitation beam. The GCaMP6s emission was detected and collected with gated GaAsP photomultiplier tubes (PMTs) for high sensitivity detection of fluorescence photons as part of the FemtO2D-Galvo scanner. The laser light transmitted through the sample was collected by the condenser lens and sent to another PMT to provide a bright-field transmission image in registration with the fluorescent images. Line scans were marked through the somata and visible dendrites and 20–250 Hz scans were performed, using ~0.2-μm pixels and an 8-12 μs dwell time. ROIs were marked manually offline based on the online marked line scans and fluorescent traces were extracted.

Optical and electrophysiological data were obtained using the software package MES (Femtonics), which also integrates the control of all hardware units in the microscope. The software automates and synchronizes the imaging signals and electrophysiological protocols. Data was extracted from the MES package to personal computers using custom-made code in MATLAB.

### Somatic current injection

To generate subthreshold depolarizations, we injected 8-12 pA for 800 ms such that the voltage almost reached the threshold of activation. A 5 ms long, 500 pA pulse was used to generate an action potential. To calculate the calcium response to the stimulations we subtracted the baseline fluorescent level 50 ms prior to the stimulations and calculated the integrated *∆F/F* over a duration 800 ms from the start time of the stimulation both for sub-and supra-threshold depolarizations.

### Spike triggered averaging (STA) and model

To generate the STA, we averaged the *∆F/F* signal around spike times. To fit the dependence of the amplitude of the STA on firing rate (Figure 6F) we used a phenomenological model. Let _τ_be the decay time constant of GCaMP. Assume a periodic neuron with an inter spike interval of duration *T*, such that *f=1/T* is the firing frequency. The decay of the GCaMP fluorescence can be modeled as 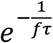. If *F*_0_ is the baseline fluorescence level and *∆F* is the amplitude of the fluorescence calcium trace visible over the baseline fluorescence, then the equation describing the decay of the GCaMP from itmaximum value to baseline value is as follows:

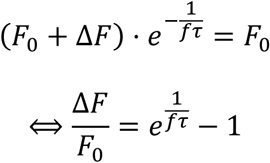

### Optogenetic stimulation

Blue light LED of the FemtO2D-Galvo scanner multiphoton system (473 nm, Femtonics) was used for full-field illumination. Light pulse trains consisted of 5 pulses at 10 Hz, each pulse lasting 1ms. Fast gated GaAsP PMTs were used to prevent saturation of the PMTs due to the LED light flashes. The PMTs were disabled 5 ms prior to LED flash and re-enabled 5 ms after the end of the light flash. To calculate the calcium response in response to the optogenetic stimulation, we subtracted the baseline fluorescence prior to the stimulations (50 ms prior to stimulation time), and compared the integrated dendritic calcium signal during a 200 ms window beginning 500 ms after evoking sub-threshold EPSPs, to the dendritic calcium signal generated by spontaneously occurring bAPs during that same time window in the same neurons.

### Immunohistochemistry

Mice were deeply anesthetized and perfused transcardially with 0.1 M phosphate buffer (PB) followed by ice-cold 4% paraformaldehyde (PFA). Coronal sections of the striatum were cut at 30 μm on a cryostat microtome (Leica CM1950) in antifreeze buffer (1:1:3 volume ratio of ethyl glycerol, glycerol, and 0.1 M PB) and stored at −20°C before further analysis. The sections were preincubated in 5% normal horse serum and 0.3% Triton X-100 in 0.1 M PB for 40 min after washing steps, and incubated overnight at 4°C with the primary antibodies [goat anti-choline acetyltransferase (ChAT), 1:100 (Millipore; RRID: AB_262156)]. On the second day, sections were incubated with fluorophore-conjugated species-specific secondary antibodies [donkey anti-goat, 1:1000 (Abcam)] for 2 h at room temperature. Brain sections were rinsed in PBS and directly cover-slipped by fluorescent mounting medium (Vectashield, Vector Laboratories). Multilabeling fluorescent immunostainings of juxtacellularly filled neurons were analyzed using a laser-scanning microscope (LSM 510 Meta, Zeiss) using 20X/0.3 NA interference contrast lens (20X zoom).

### Data and statistical analysis

#### Calculating the velocity vector of the propagation of the neuropil signal

We generated a latency map *τ*(*x*_*i*_,*y*_*j*_) of the imaged region (which is an *L*_x_ by *L*_y_ rectangle whose center is at the origin), in which the entry in each pixel (*x*_*i*_,*y*_*j*_) is the latency at which the *∆F/F* signal (averaged around all of the peak events of the spatially averaged *∆F/F* signal) peaks relative to the timing of the peak of the spatially averaged *∆F/F* signal.

We least-square fit a linear model for **τ**(*x*_*i*_,*y*_j_) which is given by τ^L^(*x*,*y*)=*p*_*x*_*x*+*p*_*y*_*y*^+^ *const*. It can be shown that direction of the propagation of the wave will be the pace vector

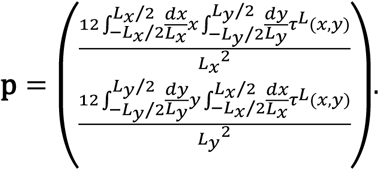

The speed of the wave will be given by ||**p**||^−1^ (to calculate the pace vector we chose the largest rectangle that could be fit into the lens region of the image).

Peak events in the spatial average were detected with a peak-finding algorithm (MATLAB) with the condition that the peak amplitude be larger than 35% *∆F/F*.

In order to further characterize the spatiotemporal structure of the spreading activity we calculated the time course of the *∆F/F* signals averaged along ten equally-spaced bars (∆x=96 µm apart) that are perpendicular to the direction of propagation. To estimate the spatial structure of the spread we rendered the time reversed version of these 10 time courses after the appropriate rescaling and shift defined by the variable transformation *x = –t||**p**||-1+n ∆x* (i.e. if the time course of the *n*^*th*^ bar’s signal is given by *f*_*n*_(*t*), then the spatial waveform at time intervals *t*_*n*_ = *n∆x*||**p**|| is given by *g*_*n*_(*x-n∆x*) = *f*_*n*_(-*x*||**p**||)).

The nonparametric two-tailed Wilcoxon signed-rank test (SRT) was used for matched samples. Null hypotheses were rejected if the *P*-value was below 0.05.

## Results

### Synchronous spatiotemporal wave-like patterns in striatal cholinergic neuropil of freely moving mice

The dorsolateral striatum (DLS) of choline acetyltransferase (ChAT)-cre mice was transfected with adeno-associated viruses (AAVs) harboring cre-dependent CGaMP6s, causing this genetically encoded calcium indicator (GECI) to express selectively in CINs (Figure 1A). Following implantation of GRIN lens in the transfected area (Figure 1B), imaging an area of approximately 600 µm by 900 µm through the lens using a miniaturized endoscope in two freely-moving mice (Figure 1C) revealed spatiotemporal fluctuations in fluorescence in the cholinergic neuropil. These fluctuations were characterized by recurring, rapid bursts of activation that permeated the entire field-of-view (FoV) and that slowly decayed (Movie 1). Embedded within the neuropil were several dozen somata of individual CINs that also exhibited substantial fluctuations in fluorescence (Figure 1D). Due to the depth-of-field of the 0.5 numerical aperture (NA) GRIN lens, the pixels from somata also contain contributions of fluorescence from the dense, space-filling cholinergic processes (Chang et al., 1982; DiFiglia, 1987; Wilson et al., 1990; Kawaguchi, 1992) traversing above and below the somata and possibly from other out-of-focus fluorescent somata. One expression of this is that the somatic fluctuations are always superimposed upon the temporal fluctuations of the surrounding pixels that contain the same contribution from the by-passing neuropil (Figure 1E, Movie 2).

**Figure 1.**
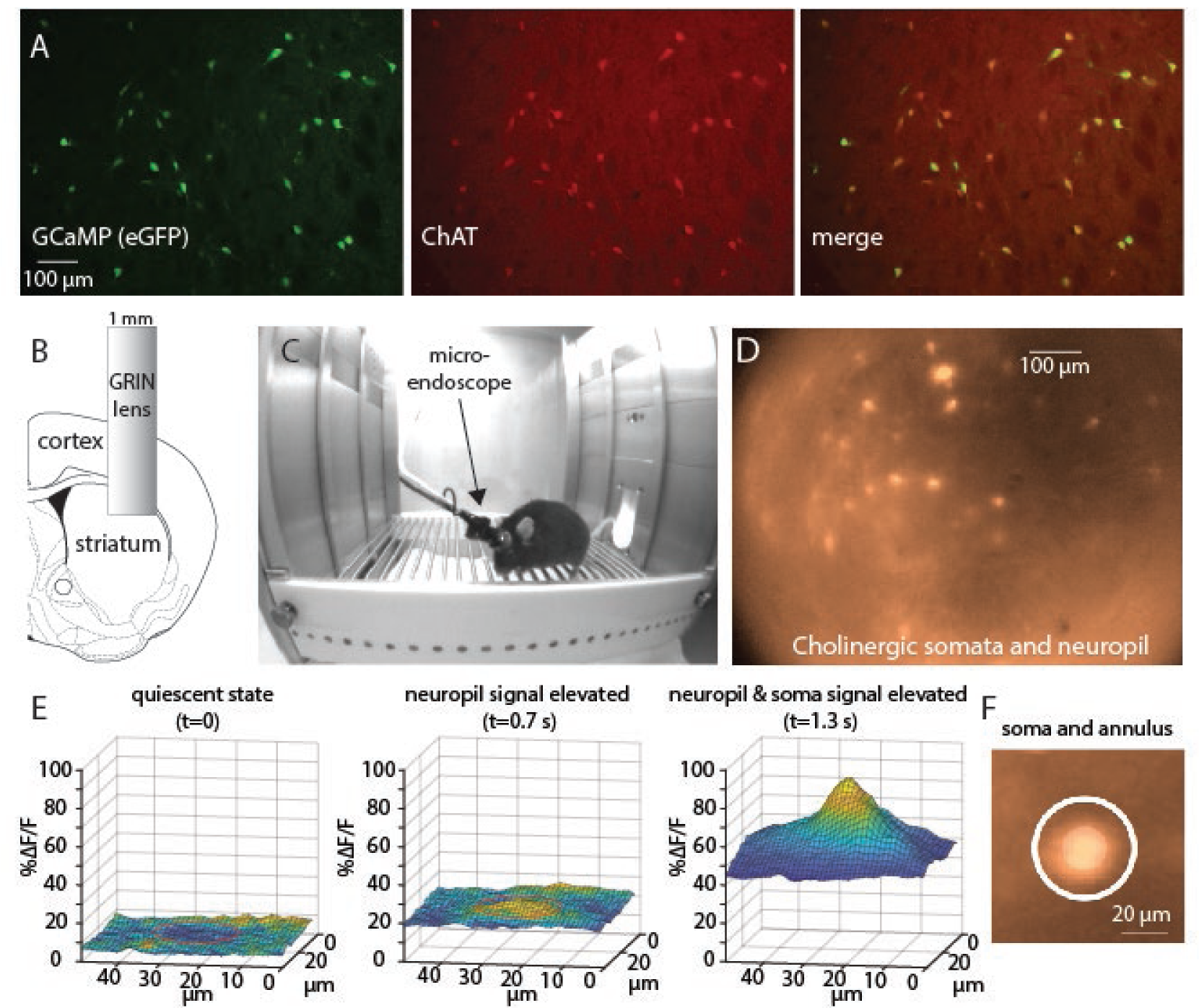
Imaging of the striatal cholinergic network in freely moving mice reveals both somatic and neuropil signals. A. Immunohistochemical analysis of dorsal striatum of choline acetyltransferase (ChAT)-cre mice stereotactically injected with adeno-associated viruses harboring floxxed GCaMP6s demonstrates its selective expression in CINs. B. A 1 mm diameter GRIN lens is implanted into dorsolateral striatum following aspiration of cortical tissue. C. Implanted mouse with a microendoscope mounted on its head moves freely in a behavior chamber. D. Image via lens in freely moving mouse reveals a GCaMP6s signal from 44 identifiable somata and from the surrounding neuropil. E. 3-D rendition of a patch of the *∆F/F* signal surrounding a soma reveals that an elevation in the neuropil signal precedes elevation of the somatic signal (region of soma indicated by a red circle). F. Illustration of the sampling of a somatic region-of-interest (central circle) and a surrounding (white) annular region.

To estimate the spatiotemporal structure of the neuropil signal and to compare it to the somatic activity, we analyzed activity in annuli surrounding each of the somata in the FoV (Figure 1F). In Figure 2A we depict a color-coded matrix of the fluctuations in fluorescence *(∆F/F*) as a function of time with each row representing an individual annulus.

**Figure 2.**
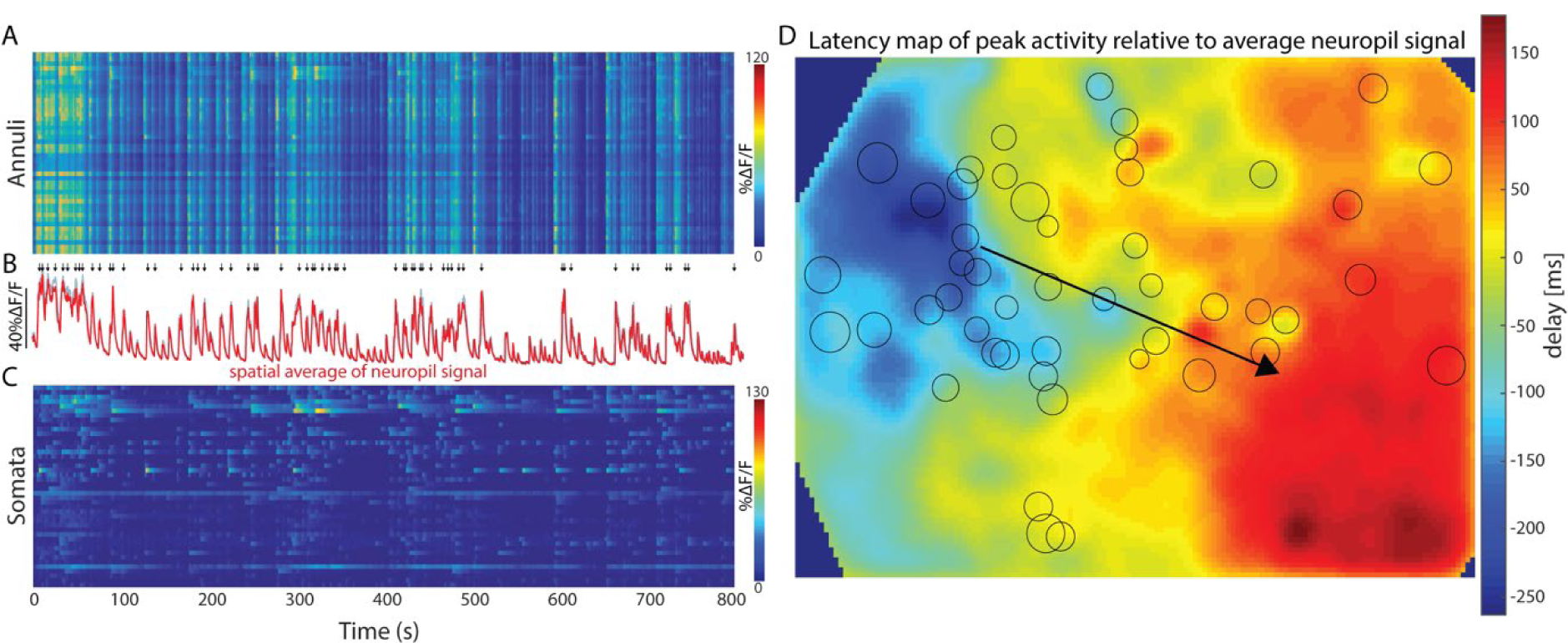
Spatiotemporal propagation of neuropil signal in dorsolateral striatum in freely moving mouse. A. Color coded matrix of activity of neuropil *∆F/F* signal sampled from the annuli surrounding 44 somata scattered in the field-of-view. Time is represented along the horizontal axis. Each row corresponds to an individual annulus. B. population average of signals from all annuli (red, using annuli associated with the somata; gray – using randomly located circular ROIs that are far from any soma). Arrows above represent peaks of strong network activation (see Materials and Methods). C. Color coded activity of the 44 somata *∆F/F* signals, after subtraction of the surrounding annular signal. D. Map of time delays of the neuropil signal peaks (in each pixel) relative to the time of peak of the average annular signal (map filtered with a 2D Gaussian filter). The arrow represents the extracted best fit direction of the velocity vector of propagation. Circles indicate the location of the annuli surrounding the individual somata.

This matrix reveals that the neuropil signal is composed of dramatic increases in *∆F/F* that are highly synchronous across the entire FoV (as is evident from the near-identical signal in the various rows of the matrix), and that decay slowly, as can be seen in the population average of all the annuli (Figure 2B, red trace). To justify the use of the annuli as a fair representation of the neuropil activity, we calculated the average signal that arises from 100 small circular ROIs randomly located within the region of the image that is devoid of somata (See Materials and Methods). The resulting average (Figure 2B, gray trace) is indistinguishable for the average annuli signal. Thus, while the annuli signals reported the global neuropil signal, the individual somata exhibited more distinct dynamics, once the signal from each annulus was subtracted from its corresponding soma (Figure 2C).

Close examination of the recurring bursts of activity (Movie 1) suggested that the neuropil signal in this mouse propagates in the plane of the image from top left to bottom right. To test this, we identified peaks of large calcium fluctuations in the average annular *∆F/F* signal (see arrows in Figure 2B). To visualize the propagation, we calculated for every pixel in the image the average latency with which its *∆F/F* signal peaked relative to the timing of the peak of the population average. A smoothened version of this map (Figure 2D) demonstrates that there is an orderly advancement of peak times from top left to bottom right. To estimate the velocity vector of this propagation, we calculated the least-square linear fit to this latency map, from which a pace vector describing the overall average propagation can be calculated (see Materials and Methods), and whose direction is depicted as an arrow along the latency map (Figure 2D).

To further investigate the spatiotemporal propagation of the neuropil activation, we defined 10 equally spaced linear ROIs perpendicular to the calculated pace vector (Figure 3A) and extracted their calcium transients. For each ROI and each detected global peak, we considered the calcium signals in a time window around the peak. We then averaged these signals across all peak events to get the average response for each linear ROI (Figure 3A). The various average traces have very similar shapes, with traces corresponding to lines at the top left of the image beginning to rise, reaching their peak and decaying earlier than those corresponding to lines at the bottom right. This is consistent with the direction of advancement depicted in the map (Figure 2D). Next, to estimate the spatial structure of the spread, we rendered the time reversed version of these 10 signals after the appropriate scaling and shift (see Materials and Methods, Figure 3B, Movie 3). The resulting ten traces correspond to different time points and show how the neuropil activation travels through the image in a wave-like manner.

**Figure 3.**
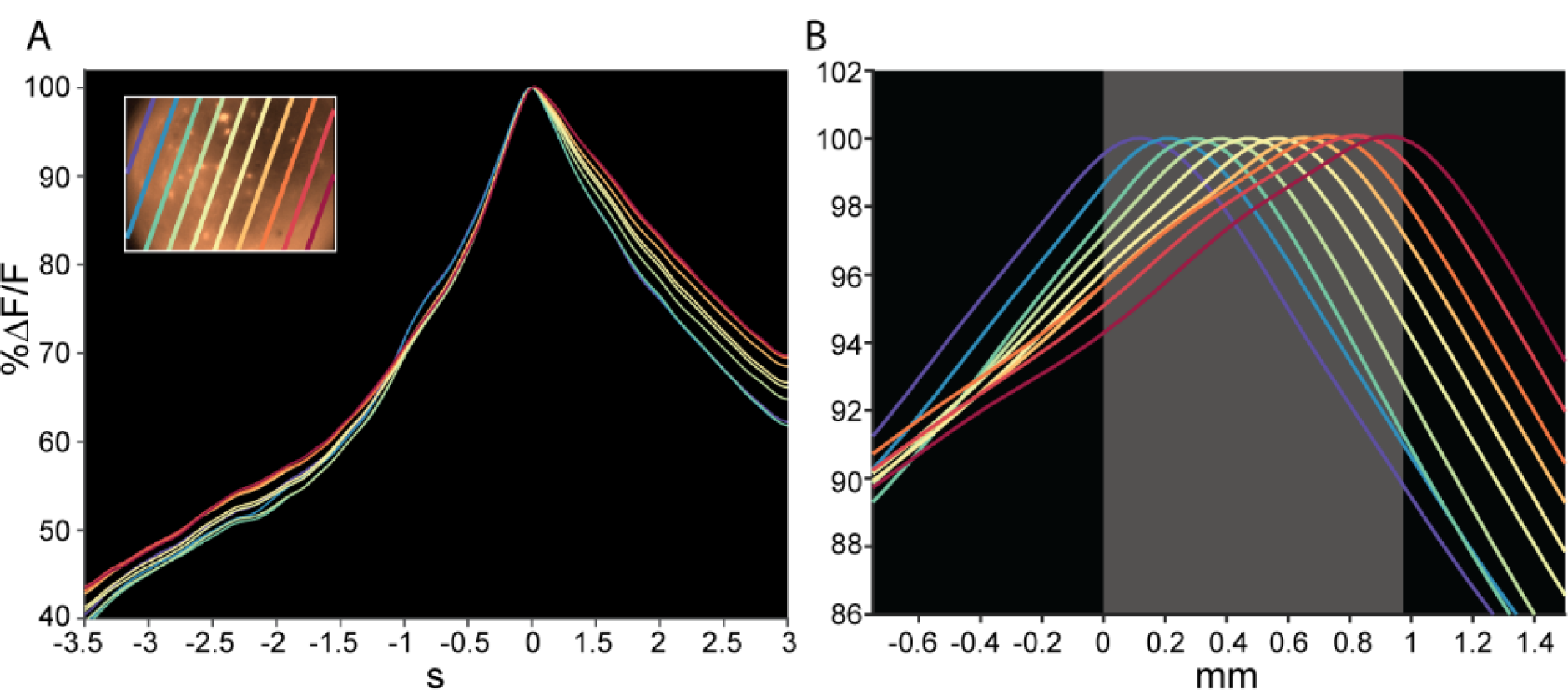
The spatiotemporal structure of activation propagation is wave-like. A. *ΔF*/*F* traces corresponding to 10 equally spaced linear ROIs perpendicular to the previously calculated pace vector. Colors of traces correspond to colors of ROIs in the inset which displays the bars over the FoV. B. Time reversed version of the 10 traces in A after appropriate scaling and shift. The various colors correspond to different time points in the propagation of the wave (see Materials and Methods). Gray area marks the FoV.

To test whether individual wave events propagated reliably in the same direction, we plotted the angular distribution of the direction of the individual maps (Figure 4A). This plot demonstrated that the individual events clustered around a single direction. The reciprocal of the magnitude of the pace vector is a measure of the wave’s speed. We found that the distribution of the speed of propagation was unimodal around 2-3 mm/s (Figure 4B). The reliability of the direction and speed of the wave propagation was also maintained in an imaging session that was a month apart. Even though the mean shifted slightly clockwise (Figure 4C), the modal lobes of the two angular distributions are still facing in the same general direction. Similarly, the modal of the distribution of the velocity also remained in the same range (Figure 4D).

**Figure 4.**
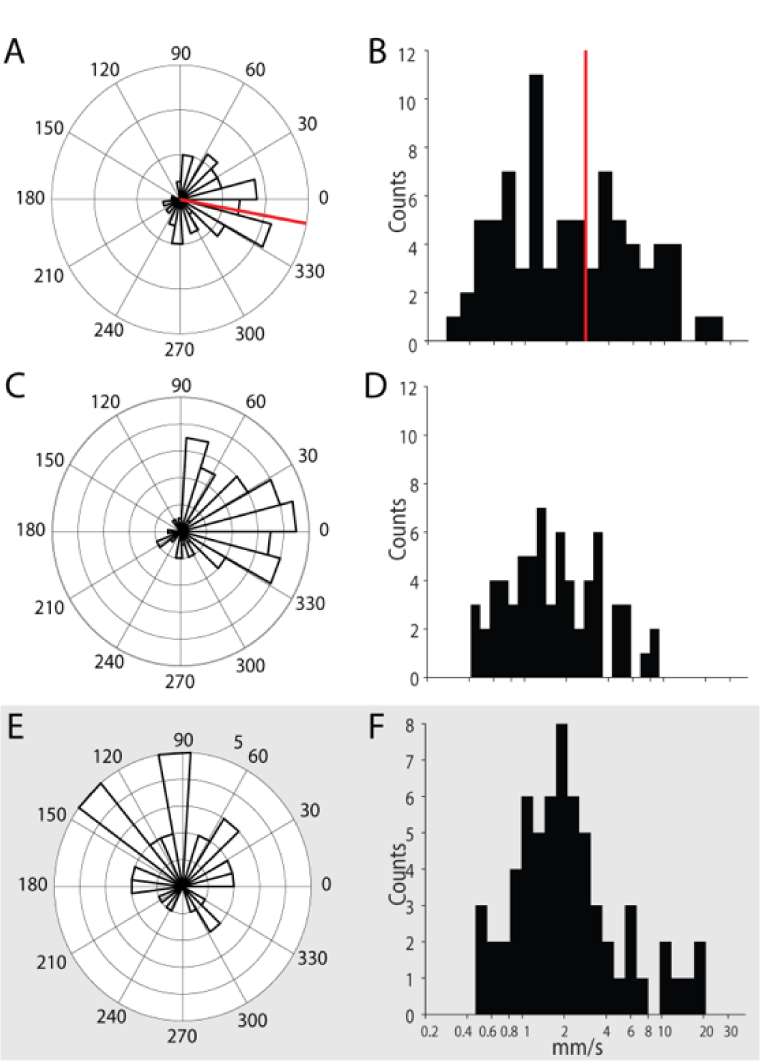
Direction and speed of neuropil traveling wave is reproducible across several weeks. A. Angular distribution of the direction of propagation of individual events that belong to the session depicted in Figure 2. B. Distribution of the speed of the individual wave events from the same session. C. Same as panel A, except from a session a month earlier. D.Same as panel B, except from a session a month earlier. E. Angular distribution of the direction of propagation of individual maps pooled from two sessions a month apart from the second mouse. F. Distribution of the speed of individual wave events pooled from the same two sessions from the second mouse. Red lines in panels A and B mark the direction and speed of the mean map depicted in Figure 2D, respectively.

An example of the same analysis from the other mouse is provided in Movie 4. In the second mouse, the direction (Figure 4E) and speed (Figure 4F) of the waves distributed in the same fashion, and remained reproducible across two sessions that were a month apart. Thus, we conclude that the cholinergic neuropil signal propagates through the striatum creating a visible wave-like progression along the dorsal surface of the striatum. Further work is needed to determine the precise directionality of these waves and their relationship to the mice’s innate behaviors.

### The neuropil calcium signal precedes the somatic signal

As seen above (Figure 1E, Movie 2), comparison of the activity of a given soma (after subtraction) and its surrounding neuropil signal (taken from the corresponding annulus) suggests that even though the activity of the ROI and annulus are quite distinct, every time a soma is activated, this activation is preceded slightly by an activation of the surrounding neuropil signal (Figure 5A). To quantify this effect, we calculated for 44 soma-annulus pairs, the event-triggered average of the annulus signal around the time of an event in its corresponding soma. The population average (across all 44 pairs) shows that each somatic event is preceded on average by a rise in the neuropil signal that begins 2 seconds earlier (Figure 5B). The neuropil also decays faster than the somatic signals (Figure 5C, median somata: 4.35 sec, median annuli: 2.32 sec, n=14 eligible pairs, *P* = 3.7×10^-4 a^, SRT). Given that the cholinergic neuropil is almost entirely intrinsic to the striatum (Mesulam et al., 1992; Contant et al., 1996; Dautan et al., 2014), it is unclear why the neuropil signal would begin prior to the somatic signals. Perhaps the neuropil signal precedes the somatic signals because the former represents input to the latter. It is possible that activation of synaptic inputs generates elevations in dendritic calcium levels that would manifest themselves as neuropil signals that precede the somatic discharge. Alternatively, perhaps the afferent cholinergic projection from the pedunculopontine nucleus to DLS (Dautan et al., 2014) was infected retrogradely by the striatal AAV injection and contributes to the fluorescence, thereby contributing to a fluorescent neuropil signal that precedes the somatic signal. However, these possibilities seem unlikely because 2 seconds is too long a latency to be accounted for simply by synaptic delays.

**Figure 5.**
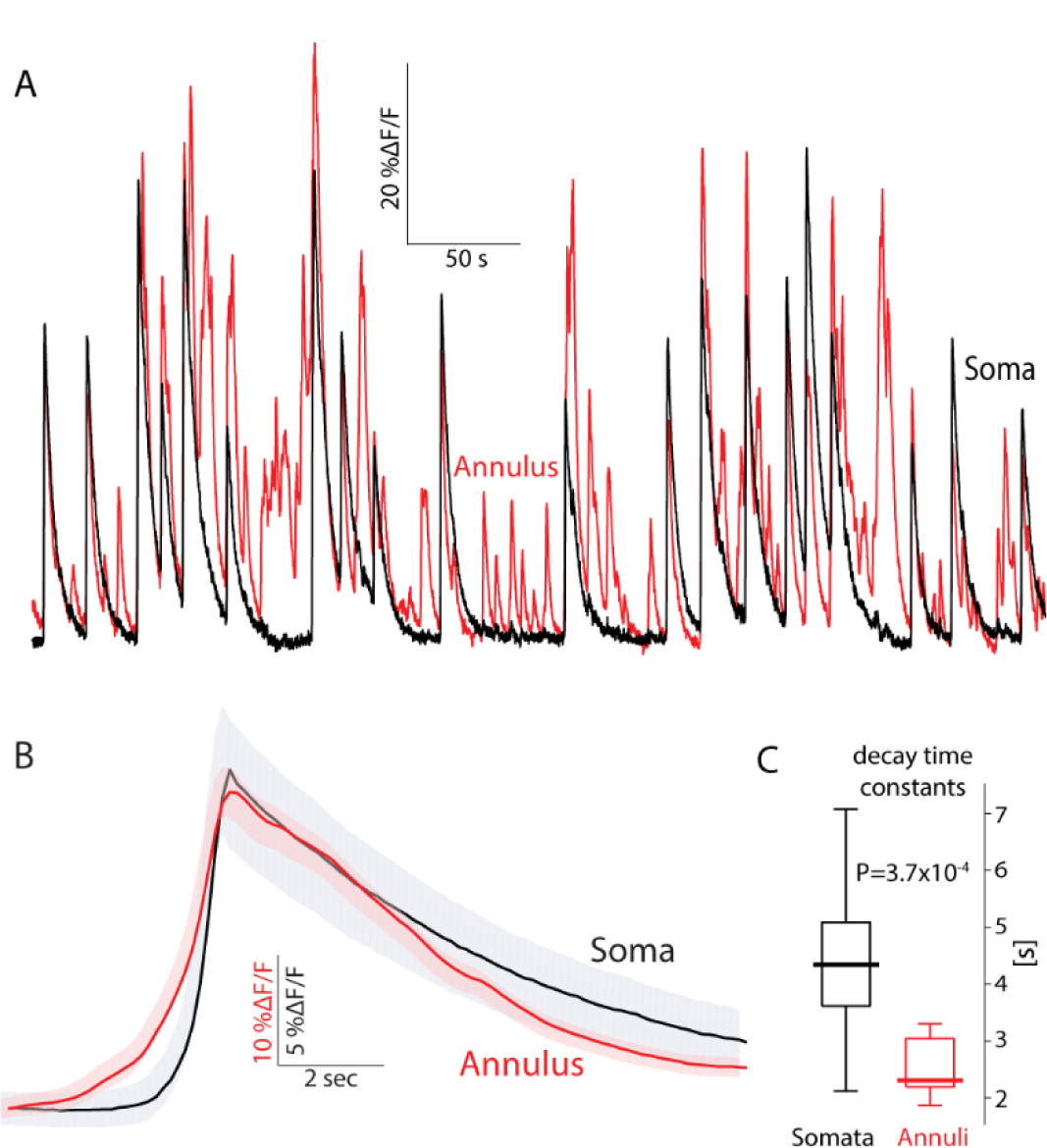
Annular (neuropil) signal precedes somatic signal in freely moving mice. A. Calcium (*∆F/F*) signal from a soma-annulus pair. B. Average calcium signal from the soma and its corresponding annulus averaged over soma-annulus pairs triggered on the somatic calcium events. Shaded areas mark the 95 percent confidence intervals. C. Boxplot of decay time constants of somatic vs. annular calcium signals. Bold line is the median and the whiskers are the 25^th^ and 75^th^ percentiles.

**Figure 6.**
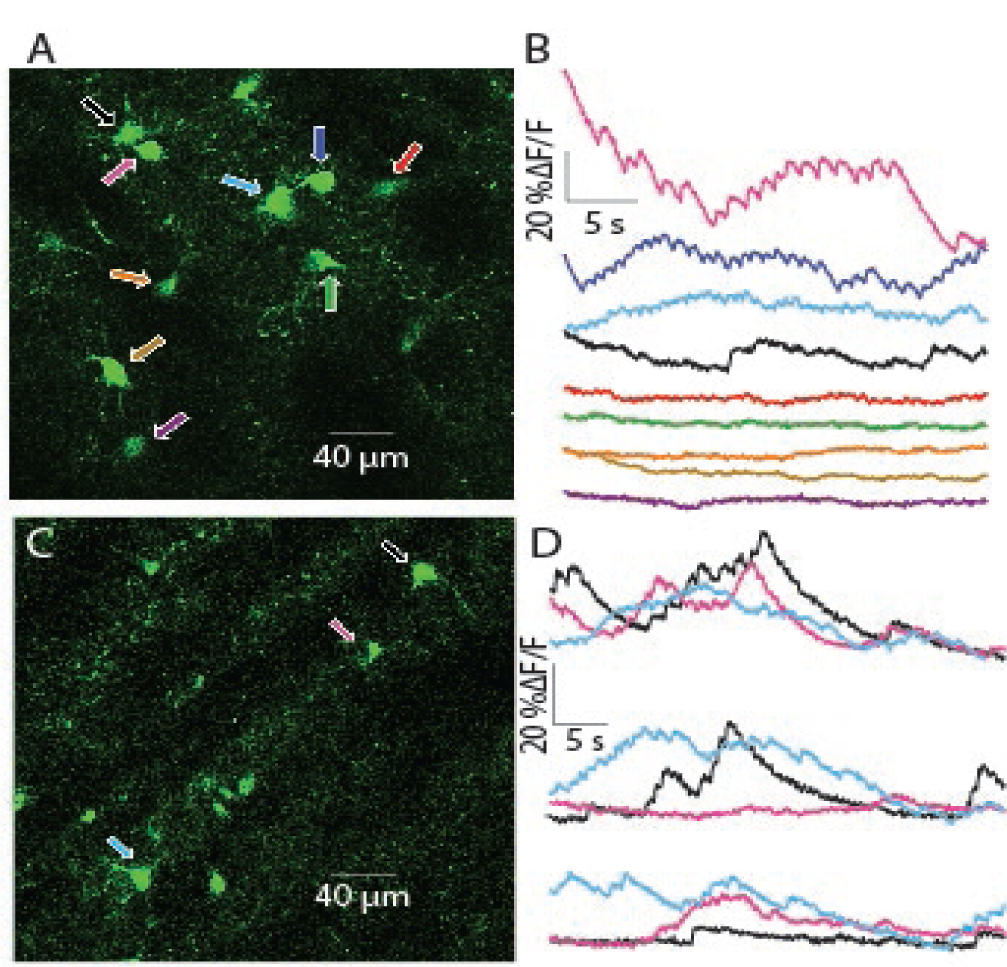
2PLSM imaging of CIN somata in acute striatal slices. A. 2PLSM image of CINs expressing GCaMP6s in an acute striatal slice. Nine individual CINs are identified and color coded with arrows. B. Color coded traces of calcium (*∆F/F*) signals from the 9 cells depicted in panel A. C. 2PLSM image from another mouse, with three identified and color-coded CINs. D. Three repetitions of calcium imaging of the 3 CINs depicted in panel C – a few minutes apart.

Several properties of the neuropil and somatic signals are therefore still in need of elucidation. How are fluctuations in somatic fluorescence signals related to the actual discharge of cholinergic interneurons? What is the physiological process that generates the neuropil signal? What is the origin of the 2 second long neuropil activity that precedes the somatic signals? These questions and the curious observation of spatiotemporal wave-like patterns in the neuropil prompted us to delve deeper into the origin of the somatic and neuropil signals generated during the collective activity of CINs.

Because CINs are autonomously active neurons (Bennett and Wilson, 1999; Surmeier et al., 2005), we reasoned that some degree of collective cholinergic activity would be preserved in acute striatal slices, and that we could image this activity in the ChAT-cre mice expressing GCaMP6s selectively in CINs. We also reasoned that using two-photon laser scanning microscopy (2PLSM) would help us discriminate between somatic and neuropil signals. Due to the miniscule depth-of-field (<2 µm FWHH) of 2PLSM, the contamination of somatic imaging by out-of-focus neuropil would be negligible. Additionally, 2PLSM allows imaging of individual dendrites which are presumably a major contributor to the neuropil signal, and study precisely under what circumstances they give rise to GCaMP6s signals. By combining targeted patch clamp recordings with 2PLSM imaging we could address mechanistic questions regarding the relationship of the somatodendritic calcium signal and the electrophysiological activity of the CINs.

### Somatic GCaMP6s signals exhibit diverse temporal patterns of activity *ex vivo*

2PLSM imaging of acute striatal slices demonstrated that neuropil expressing fluorescent GCaMP6s could be observed. Moreover, the neuropil can be resolved as dendritic (and perhaps axonal) processes. The striations were devoid of GCaMP6s, in line with their known composition of non-cholinergic afferent fibers (Wilson, 2004). Embedded within the fluorescent neuropil processes several (up to 10 per experiment) CIN somata could be visualized. In the example depicted in Figure 6A, we were able to manually mark 9 distinct somata for imaging and extract traces of *∆F/F* that exhibited autonomous calcium fluctuations in the transfected neurons (Figure 6B). In some traces, calcium transients that appear to correspond to individual action potentials could be observed (top 3 traces). Collectively, these traces demonstrated that the temporal structure of the fluctuations varied considerably among the somata (Figure 6B), with some cells exhibiting large fluctuations in *∆F/F* while others exhibited relatively uneventful traces (bottom 5 traces). Interestingly, the nature of the calcium signal in individual neurons could vary considerably with time. For example, one CIN (marked by the pink arrow in Figure 6C) exhibited large calcium fluctuations in the initial trace (Figure 6D, upper pink trace). A few minutes later, these fluctuations died down (middle pink trace). Finally, after a few more minutes elapsed, large fluctuations resumed (bottom pink trace). Such findings are consistent with previous studies that showed that individual CINs in acute slices can change their firing pattern from regular to irregular to bursting discharge (Bennett and Wilson, 1999).

It is easy to make a convincing case that the GCaMP6s signal represents autonomous discharge when transients associated with individual spikes or large fluctuations in *∆F/F* are observed. However, what can be said for the cells that exhibit a relatively flat and unchanging profile of fluorescence?

### Limitations in interpreting somatic GCaMP6s signal from autonomously firing CINs

To address this question, we used targeted-patch of CINs in order to characterize the firing patterns underlying the calcium transients (Figure 7A). We then aligned the trace from the somatic region of a cell with its corresponding electrophysiological recording. When a CIN discharged spontaneously in a bursty fashion (Figure 7B, blue), calcium transients similar to those observed without a patch electrode (Figure 6) were visible, and it was possible to clearly associate the electrophysiological recordings and the large calcium transients. Proximal dendrites exhibited an even clearer relationship to spiking (Figure 7B, pink), due to the larger surface-to-volume ratio (see below). In CINs that fired regularly (Figure 7C), fluorescence transients associated with single actions potentials could still be identified, as long as the firing rate was sufficiently low (Figure7D, top). In this case, estimation of the spike triggered average (STA) of the calcium signal revealed a large *∆F/F* waveform that rose after the action potential occurred and slowly decayed (Figure 7E, top). When the same CIN discharged faster, it was no longer possible to unequivocally discern individual transients (Figure 7D, bottom). Nevertheless, the STA revealed a *∆F/F* waveform that is time-locked to the action potential (Figure 7E, bottom), albeit a significantly smaller one than that observed during slower discharge (Figure 7E, top). This finding suggested the tetanic-like fusing of the individual calcium transients is responsible for the decrease of the STA amplitude. If tetanic fusion of the *∆F/F* signal occurs at firing rates as low as 2-2.5 spikes/s, this effect places strong constraints on the ability to read out the discharge patterns underlying the calcium transients when using GCaMP6s.

**Figure 7.**
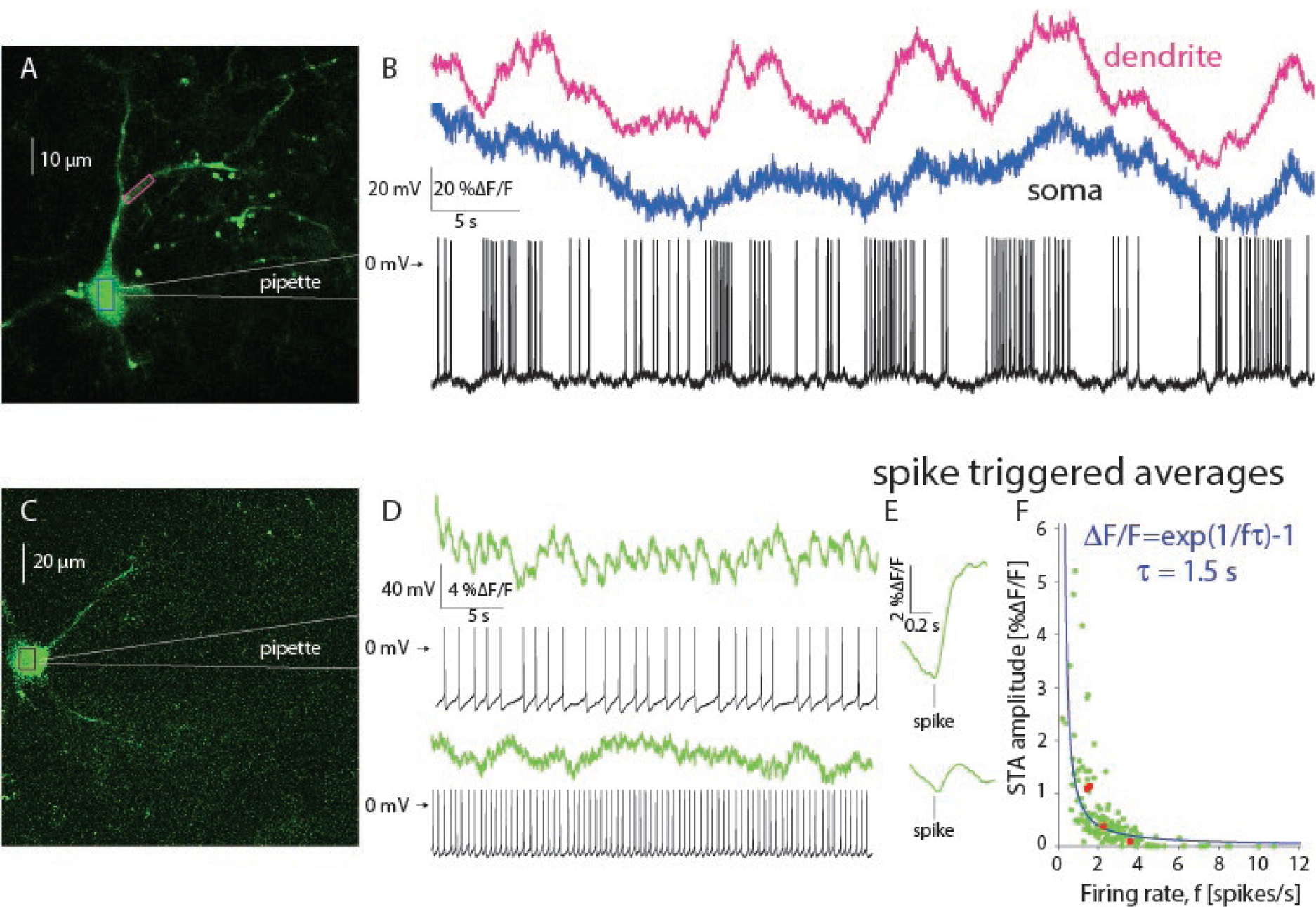
Estimation of autonomous discharge of CINs using 2PLSM GCaMP6 imaging is limited to bursting or to slow regular firing neurons. A. 2PLSM imaging of somatic and dendritic calcium (*∆F/F*) signals in conjunction with electrophysiological recording from GCaMP6s-expressing CIN. B. Somatic (blue) and dendritic (pink) *∆F/F* signals and membrane potential of CIN depicted in panel A. C. 2PLSM somatic imaging and electrophysiological recording from another CIN. D. Somatic imaging and corresponding electrophysiological recording from CIN firing autonomously at a lower rate (top) or driven to fire faster at a higher rate (bottom). E. Spike-triggered average (STA) of somatic *∆F/F* signal (from panel D) for slow (top) and fast (bottom) discharge. F. Amplitude of STA vs. the firing rate of regularly firing CINs. Solid blue line: fit of phenomenological model (see text).

To quantify this effect, we characterized the relationship between firing rate of the CINs and the amplitude of the STA of the *∆F/F* trace associated with that firing rate. Figure 7F depicts a scatter plot of the STA amplitude as a function of firing rate for *n=25* non-bursting CINs (from *N=15* mice), where each CIN is represented by at least three points corresponding to different firing rates (e.g., red dots represent 4 measurements from the same CIN). The resulting plot (Figure 7F) shows that the STA calcium transient amplitudes decrease as the firing rate increases. When the firing rate is too high it becomes impossible to detect the underlying spike times from the fluorescence trace alone, because the average transient is too small to discern against a noisy background. An example for such a case is shown in Figure 7D, bottom, where the neuron fires at ~2-2.5 Hz. Individual action potentials can no longer be discerned from the imaging trace alone even though the estimation of the STA reveals a visible waveform.

The dependence of the size of the STA on the firing rate can be captured by a simple phenomenological model (see Materials and Methods) wherein the observed calcium dynamics are simply a sum of the calcium fluorescence transients elicited by the individual action potentials. If we further assume for simplicity that the transient is exponentially shaped (with a decay time constant τ) and that the firing is perfectly periodic, we can derive an expression for the amplitude of the STA as a function of the firing rate that depends on the single parameter τ. The value of *τ*for the fit (blue line in Figure 7F) is 1.5 seconds. The main determinant of this decay is the decay kinetics of GCaMP6s which is on the order of one second (Chen et al., 2013). Additionally, imaging large compartments such as somata further slows the decay kinetics (Goldberg et al., 2009).

### Capacity to detect individual spikes using wide field calcium imaging is diminished

From the empirical results and the phenomenological model, it is evident that the faster the neurons fire, the smaller the size of the STA waveform. In other words, the slower the decay kinetics of the indicator the harder it is to discern individual action potentials in the calcium signals. These data demonstrate that the use of GCaMP6s, constrains the ability to detect individual calcium transients if the neuron fires faster than 2-3 spikes/s. But what resolving power is GCaMP6s expected to have under the one-photon conditions employed in the endoscopic imaging? Unlike the sensitivity of 2PLSM imaging, the endoscopic data collected from the awake behaving mouse did not seem to exhibit calcium transients corresponding to individual action potentials. One possibility is that the much slower sampling rate of the endoscope – which smears STAs (Teagarden et al., 2008) – and its low NA (0.5) GRIN lens objective further diminishes the ability to discern individual action potentials. To test this possibility, under more controlled conditions in conjunction with electrophysiological recordings, we conducted wide-field calcium imaging of CINs expressing GCaMP6s in acute slices under conditions that recreate the endoscope’s specification, by using an EM-CCD at a sampling rate of 5-10 Hz via a 20X/0.5 NA objective.

The wide-field imaging revealed a diffuse neuropil signal, similar to that seen *in vivo* (Figure 8A). As with the 2PLSM imaging experiments, we were able to discern multiple CINs per slice (Figure 8A), and the calcium transients associated with individual neurons were variable, with some exhibiting large and slow fluctuations while others exhibiting flat traces (Figure 8B). When we combined electrophysiological recordings with the imaging, it was possible to observe gross changes in firing rate in the *∆F/F* signal (Figure 8C). However, when we focused on examples where the discharge was relatively regular, it was impossible to observe fluorescence transients associated with individual spikes. Furthermore, the STAs in these cases were indistinguishable from a flat line (*n=9* neurons from *N=8* mice, data not shown).

**Figure 8.**
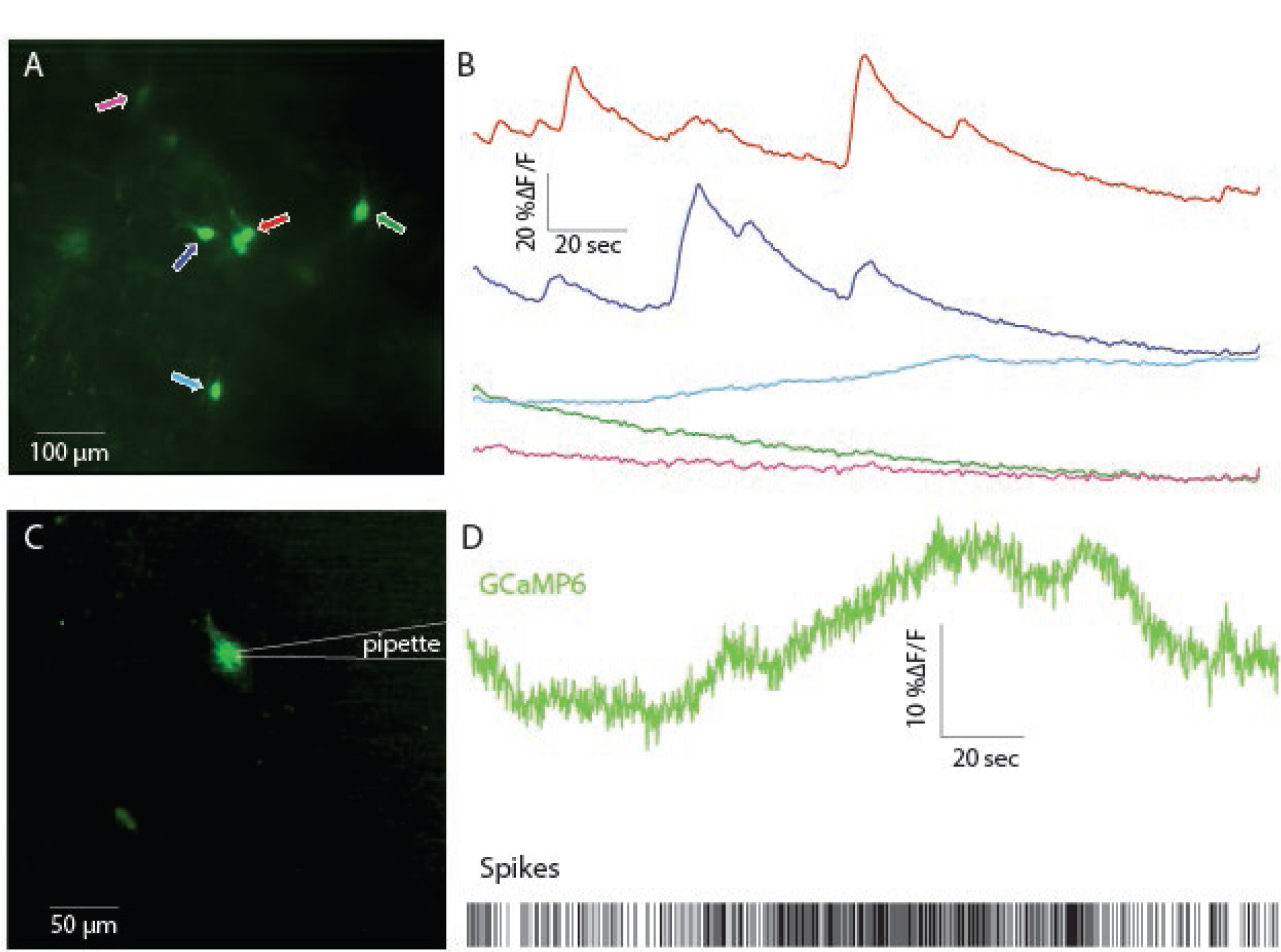
Wide-field one-photon imaging of CIN somata in acute striatal slices. A. Wide-field image of CINs expressing GCaMP6s in an acute striatal slice. Five individual CINs are identified and color coded with arrows. B. Calcium (*∆F/F*) signal from the 5 cells depicted in panel A. C. Wide-field image from another cell recorded in cell attached mode. D. Calcium (*∆F/F*) in conjunction with spike raster of the CIN depicted in panel C.

Our conclusion is that a sufficiently slow firing rate is required in order to discern calcium transients from individual spikes, and that – in our hands – this is achieved reliably only using 2PLSM. Wide field imaging with a slow camera and a low NA objective will generate GCaMP6s signals, that follow changes in firing rate, but from which individual spikes are not detectable. Moreover, when observing a calcium trace with only small fluctuations it is difficult to determine whether the cell is firing fast or not firing at all. The conclusion is that with the use of endoscopes one can only observe substantial fluctuations in firing rate. Moreover, because constant fluorescent signals are usually included in the baseline value (*F*_0_), any relatively regular activity of CINs will generate a basal fluorescence that will be removed when calculating the *∆F/F* signal, and will therefore remain undetected.

### Origin of the neuropil signal

The previous analysis suggests that the fluctuating somatic GCaMP6s signals observed via the endoscope extract only robust changes in the firing rate of CINs. We next turn to consider what neural activity gives rise to the cholinergic neuropil signal observed in endoscopy. Here again we employ 2PLSM in acute striatal slices with GCaMP6s-expressing CINs. As mentioned above, distinct dendritic processes are observed with 2PLSM throughout the transfected regions in the striatal slice. As these processes are ubiquitous, and no other cell types express the GECI, we hypothesized that CIN dendrites contribute to the blurred neuropil signal visualized with the endoscopes and the wide field calcium imaging (Figure 1D, movies 1 & 2).

Because of the large, global fluctuations of the neuropil signal in the endoscopic imaging, we set out to determine their source. Previous studies of calcium signals in CINs from acute rodent slices using inorganic membrane impermeable dyes (Bennett et al., 2000; Goldberg et al., 2009; Tanimura et al., 2016) demonstrated that bAPs and even long back-propagating subthreshold depolarizations (bSDs), can open dendritic voltage-activated calcium channels (Chen et al., 2013), and generate visible dendritic calcium changes. These are certainly candidate mechanisms to generate dendritic GCaMP6s signals. However, because the neuropil signal preceded the somatic signals in the endoscopic data, we considered another possibility, that activation of afferent glutamatergic inputs could introduce sufficient calcium influx via NMDA and calcium permeable AMPA receptors, both of which are expressed in CINs (Consolo et al., 1996; Kosillo et al., 2016; Aceves Buendia et al., 2017), to generate a visible calcium signal. In order to determine which of these three scenarios – namely bAPs, bSDs or activation of ionotropic glutamate receptors – is responsible for the dendritic signals, we compared the size of the GCaMP6s *∆F/F* signals generated by each of these physiological processes.

### Back-propagating action potentials are a major contributor to the neuropil signal

To determine whether bAPs or bSDs are the main contributors to the dendritic calcium signal, we target-patched CINs, silenced them with a hyperpolarizing holding current, and injected sub-or supra-threshold depolarization currents while measuring the resulting calcium transient at a proximal dendrite. Comparison of the size of the transients evoked by back-propagating sub-and supra-threshold depolarization revealed that bAPs generated a significantly larger calcium signal in dendrites, that were quantified by the integrated fluorescence beneath the *∆F/F* curve for a period of 800 ms (median bAP integrated fluorescence: 0.6875, median bSD integrated fluorescence: 0.048, *P* = 7.8×10^-3 b^, SRT, data not shown). This result suggests that of the first two possibilities, bAPs are the main source of the GCaMP6s neuropil signal, whereas the contribution of sub-threshold calcium entry is negligible.

In order to test whether synchronous activation of glutamatergic inputs could also generate visible calcium transients in CIN dendrites, we injected mice with AAVs harboring channelrhodopsin-2 (ChR2) with a ubiquitous promoter into the thalamic parafascicular nucleus (PfN), which gives rise to the dominant glutamatergic input to CINs (Lapper and Bolam, 1992; Ding et al., 2010; Threlfell et al., 2012; Aceves Buendia et al., 2017). Thus, in acute slices from these animals, CINs expressed GCaMP6s while thalamic fibers expressed ChR2. Two-to-three weeks after transfection, we target-patched CINs (Figure 9A) and evoked excitatory synaptic potentials (EPSPs) in the patched CIN using full-field 470 nm LED illumination (Figure 9B). We compared the integrated dendritic calcium signal in response to sub-threshold EPSPs to the dendritic calcium signal generated by spontaneously occurring bAPs in the same cells. The calcium transient generated by spontaneous bAPs was significantly larger than the transient that corresponded to the sub-threshold EPSPs (Figure 9C, median bAP transient integrated fluorescence: 0.2803, median EPSP transient integrated fluorescence: 0.0382, *P* = 0.0195^c^, SRT. The values for subthreshold current pulses are shown, as well, for comparison).

**Figure 9.**
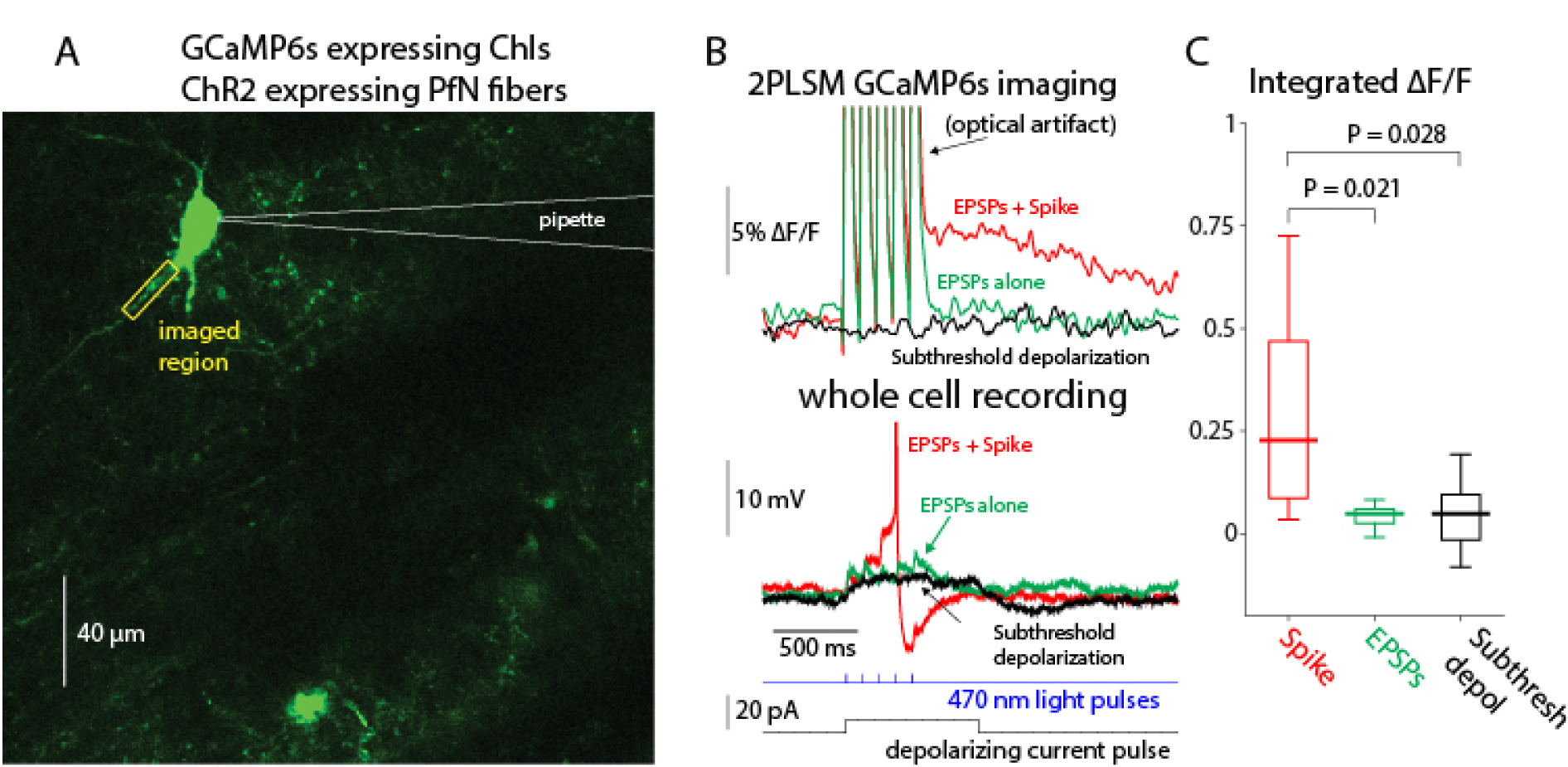
Dendritic GCaMP6 signals detectable using 2PLSM imaging are limited to back propagating action potentials (bAPs). A. 2PLSM imaging of somatic and dendritic calcium (*∆F/F*) signals in conjunction with electrophysiological recording from GCaMP6s-expressing CIN surrounded by parafascicular nucleus (PfN) fibers expressing ChR2 (not shown). B. Calcium (*∆F/F*) signals in response to optogenetic activation of PfN synapses elicits either a subthreshold (green) or suprathreshold (red) synaptic response (bottom) in CIN depicted in panel A. Only the spiking response elicits a detectable *∆F/F* signal. Subthreshold depolarization (in another cell) does not elicit a detectable response (compare to Goldberg et al. 2009). C. Distribution of integrated *∆F/F* response in response to spontaneous bAPs, subthreshold EPSPs and subthreshold depolarization.

**Figure 10.**
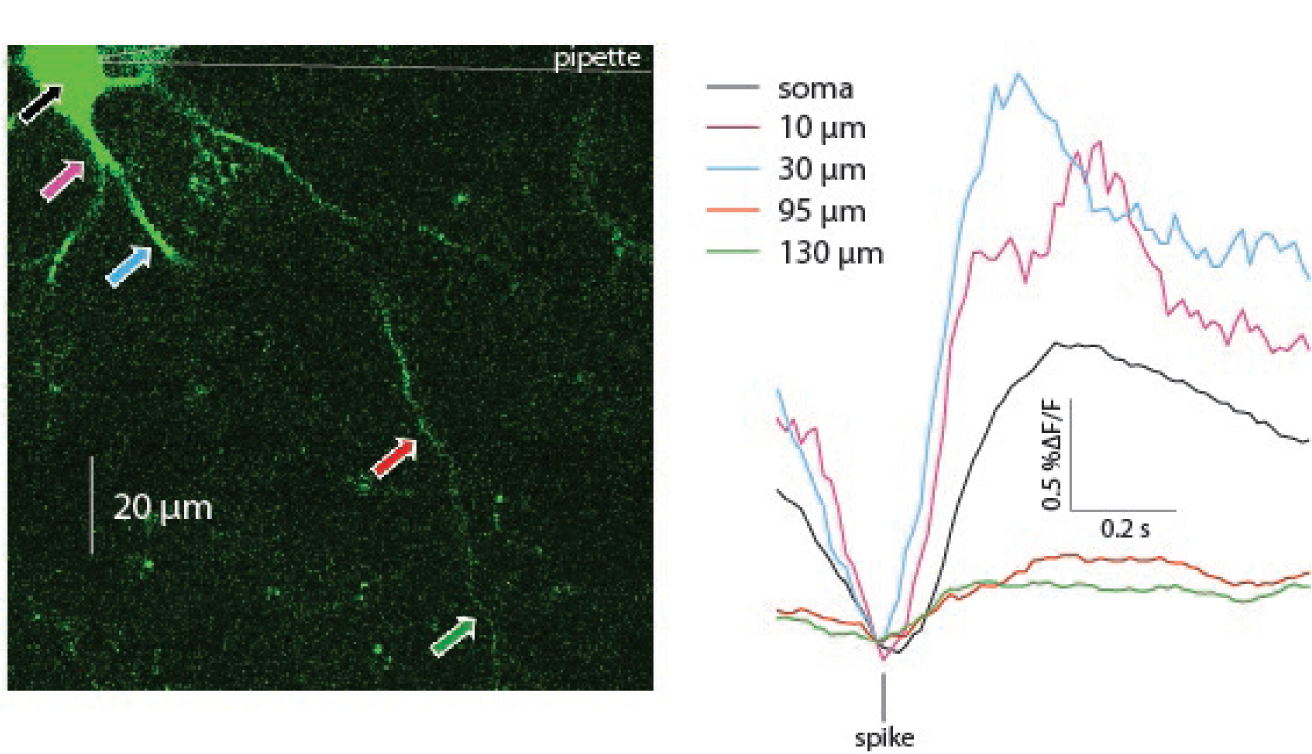
Spike triggered averages of calcium transients elicited by bAPs at various distances from the soma. 2PLSM line scans performed at various linear distances from the soma of a GCaMP6s expressing CIN.

Because the slice is filled with many fluorescent processes it was only possible to reliably identify the proximal dendrites belonging to the patch-clamped CIN. Thus, strictly speaking the previous analysis relates to calcium transients in proximal dendrites. Nevertheless, it was occasionally possible to observe bAPs infiltrating distal regions, up to 150 µm from the soma (Figure 10), as reported previously (Tanimura et al., 2016). We conclude that only bAPs generate a visible dendritic calcium signal. Indeed, as expected from surface-to-volume ratio considerations, and as has been shown previously with inorganic calcium indicators (Bennett et al., 2000; Goldberg et al., 2009) the ongoing GCaMP6s *∆F/F* fluctuations are significantly larger in the dendrites than in the soma (Figure 7), as observed by comparing STAs of dendritic transients to STAs of somatic transients (*n=23* cells from *N=13* mice, *P* = 7.35×10^−4 d^, SRT)(Figure 11).

**Figure 11.**
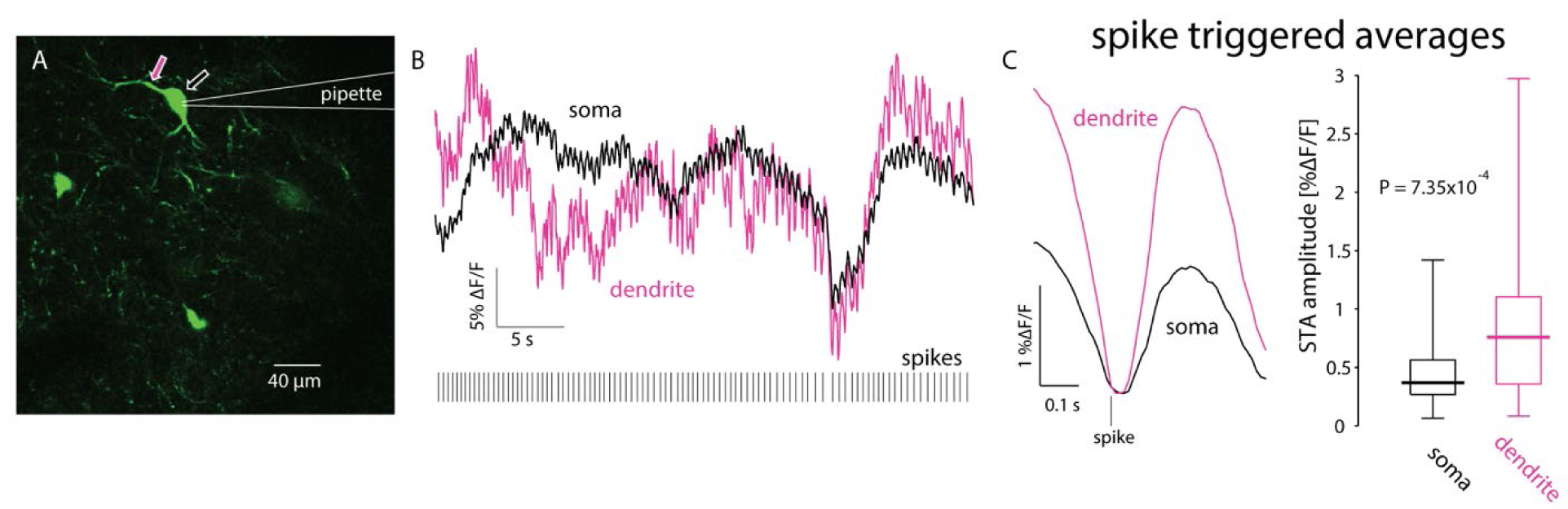
Dendritic calcium transients are larger than somatic ones. A. 2PLSM imaging of somatic and dendritic calcium *(∆F/F*) signals in conjunction with electrophysiological recording from GCaMP6s-expressing CIN. B. Calcium (*∆F/F*) responses in soma (black) and proximal dendrite (pink) during spontaneous discharge (bottom). C. STA of proximal dendritic (pink) calcium signals is larger than the somatic (black) signal. Right: paired comparison of the amplitude of somatic and dendritic STAs.

### The neuropil signal represents a mean-field readout of the entire CIN network

In light of the above experiments, it is likely that the background activity seen in the endoscopic *in vivo* imaging results from bAPs arising from CINs scattered throughout the striatum whose somata may not be visible but whose dendrites are within the FoV. While axons are not readily visible in these experiments (Bennett et al., 2000; Goldberg et al., 2009), we cannot rule out the APs propagating along the CINs’ axonal arbors also contribute to the background activity. Whenever a GCaMP-expressing CIN fires, its entire dendritic (and possibly axonal) arbor lights up. Because the axonal and dendritic arbors of CINs overlap and cover the whole striatum (Chang et al., 1982; DiFiglia, 1987; Wilson et al., 1990; Kawaguchi, 1992), a pixel whose depth of field is tens of microns integrates the fluorescent signals of bAPs from dendrites of many neurons. Nearby pixels will integrate similar signals and will therefore be highly correlated. Because this correlated signal is the sum of the activity of many cells it represents a “global mode” of CINs activity. Because the global mode arises from the bAPs in CINs, it can be considered a read-out of the “mean-field” of the recurrent network activity of the entire CINs population.

If the neuropil signal is a mean-field readout of the recurrent CIN network, then it is clear why on average the peak of the somatic signals coincides with the peak of the neuropil signal. What then gives rise to the 2 second rise in the population response that precedes the peak response of individual CIN soma? The most likely explanation is that the population (neuropil) signal represents the gradual recruitment of many CINs. Consequently, it is slower to rise than the signal from each individual neuron (as seen in the somatic signals). Additionally, when sampling a single neuron from a large population that is being recruited, it is expected that the onset of the signal from the single neurons will follow the population average. This is particularly true in our case where we sample neurons that are on the dorsal surface of the striatum, and where it is likely that the activation begins in the more ventral bulk of the striatum. We therefore conclude that the neuropil signal represents bursts of concurrent neuronal discharge of the striatal cholinergic network as a whole. This signal can be considered a mean-field cholinergic signal that is complementary to the signal recorded from individual CINs. Additionally, our data reveal that this signal propagates spatially – exhibiting wave-like patterns – throughout the striatum. The 2 second long recruitment process means that with a typical speed of 2 mm/s the wave covers most of the striatum. Nevertheless, it is quite remarkable given the time scales of the animals’ innate behaviors. Future work is necessary to determine the behavioral correlates of this propagation.

**Table 1.** Statistical Table

|  | **Data**<br>**Structure** | **Type of Test** | **Power** |
| --- | --- | --- | --- |
| a | Ranks | Two-tailed Wilcoxon signed-rank test for paired replicates | $3.7 \times 10^{-4}$ |
| b | Ranks | Two-tailed Wilcoxon signed-rank test for paired replicates | $7.8 \times 10^{-3}$ |
| c | Ranks | Two-tailed Wilcoxon signed-rank test for paired replicates | 0.0195 |
| d | Ranks | Two-tailed Wilcoxon signed-rank test for paired replicates | $7.35 \times 10^{-4}$ |
| **Table 1. Statistical Table** |  |  |  |

### Discussion

The combination of GECIs and microendoscopic imaging provides an exciting opportunity for studying the collective dynamics of striatal CINs in freely moving animals. CINs are autonomously active (Bennett and Wilson, 1999; Surmeier et al., 2005) and have extensive axonal and dendritic arbors (Chang et al., 1982; DiFiglia, 1987; Wilson et al., 1990; Kawaguchi, 1992). These properties generate a significant neuropil calcium signal. Our investigation of this signal uncovered a synchronous spatiotemporal wave-like pattern of activation in striatal cholinergic neuropil. Our analysis suggests that bAPs are major contributors to the neural activity giving rise to the background signal seen in endoscopy. Finally, we show that the neuropil signal acts as an order parameter representing the striatal CIN network as whole and is, in this sense, similar to LFP.

Performing endoscopic calcium imaging of pacemaking neurons raises several issues that require careful consideration. First, the proper method of analyzing calcium signals using *∆F/F* as a proxy for changes in calcium concentrations does not allow the detection of baseline spiking activity exhibited by regularly firing pacemakers. We combined calcium imaging and slice electrophysiology to demonstrate that 2PLSM calcium imaging using GCaMP6s only reveals individual spikes in pacemakers if their discharge rate is sufficiently low (~2 spikes/s). This is due to the relatively long decay time constant of GCaMP6s (Chen et al., 2013) that does not allow for the sufficient decay of calcium signals between consecutive spikes. Somatic calcium dynamics may introduce an additional delay, further hindering the detection of single action potentials. In the case of endoscopic imaging, the low sampling rate and low NA associated with miniaturized endoscopes add a major constraint, which makes it difficult to detect individual spikes even with averaging. Nevertheless, endoscopic calcium imaging has critical advantages over electrophysiological recordings -in particular, the ability to detect spatial patterns in the activity of a molecularly identified neural population.

We exploited this advantage to demonstrate a spatiotemporal structure in the recruitment of striatal cholinergic neuropil. The temporal recruitment is manifested in the earlier rise of the integrated neuropil signal compared to the somatic signals of individual neurons. The spatial organization of neuropil recruitment takes the form of a wave-like activity, with a reproducible direction of propagation. The identification of a systematic direction (e.g., between DLS and dorsomedial striatum (DMS)) requires future work, as it is likely to depend on the behavioral context and state of the animal. One possibility is that CINs in the DLS, believed to be involved in the execution of habitual behaviors, will be activated first when the animal is executing a familiar action, while the CINs of the DMS may be activated first during exploration and goal-directed behaviors (Yin et al., 2006; Ikemoto, 2007). It may also be that the robustness of the spatial patterns depends on the level of proficiency of acquired behaviors, with clearer wave-like activity forming during acquisition of an association crucial for the behavior that subsides later on. Finally, it could be that wave-like activity is also present in the neuropil signal of other striatal neurons. These questions will be addressed through future work, partly by studying the relationship of the collective activity of cholinergic somata and neuropil to self-initiated and learned behaviors.

Our results suggest that the background neuropil signal seen in endoscopic *in vivo* calcium imaging arises primarily from the integrated fluorescent signal of bAPs from many CINs dispersed throughout the striatum (we cannot entirely rule out that subthreshold or synaptically driven calcium influx contribute to the signal, but their contribution is negligible in comparison to suprathreshold calcium influx). The fact that the decay time of the neuropil signal is shorter than that of the somatic signals (Figure 5C), supports the interpretation that it arises primarily from dendrites (and/or axons) because their surface-to-volume ratio is higher. The higher ratio speeds up the decay constant of the axodendritic fluorescent signals relative to somatic ones (Goldberg et al., 2009). Thus, the cholinergic neuropil signal acts as a mean-field readout of the recurrent activity of the network of striatal CINs. This is reminiscent of the LFP signal – a once neglected signal that is now extensively studied and believed to represent integrated synaptic activity (Creutzfeldt et al., 1966; Eggermont and Smith, 1995; Bedard et al., 2004; Goldberg et al., 2004). We show that fluctuations in neuropil fluorescence are, like fluctuations in the LFP signal (Eckhorn and Obermueller, 1993; Destexhe et al., 1999), highly correlated in space. In addition, somatic calcium events are preceded by neuropil calcium events. This again resembles the LFP signal, which coincides with increases in instantaneous firing rate (Lass, 1968; Donoghue et al., 1998; Destexhe et al., 1999). Thus, the background neuropil signal is a signal with physiological significance.

One of the oldest ideas in striatal physiology is that a reciprocal balance between striatal acetylcholine and dopamine levels is necessary for the normal functioning of the striatum (Lehmann and Langer, 1983). Dopamine is a critical neuromodulator mediating both the motivation to engage in a current behavior and the learning of future behaviors (Schultz et al., 1993; Cagniard et al., 2006; Shiner et al., 2012; Berke, 2018). Recent studies show that dopamine signals arising from the midbrain and the striatum are dissociated and may be involved in distinct dopaminergic functions (Howe and Dombeck, 2016; Kim et al., 2016; Parker et al., 2016). This requires dopamine recipient circuits to actively switch the way that they interpret dopamine, and striatal CINs have been proposed as the natural candidates for regulating this switch (Morris et al., 2004; Yamanaka et al., 2018). Thus, the spatial propagation of cholinergic activation might also elicit hot spots and even traveling waves of dopamine concentrations in the striatum. This intriguing idea could be tested in future studies using new optical probes that enable the visualization of local changes in the concentration of dopamine, as well as other neurotransmitters (Patriarchi et al., 2018).

## Author Contributions

RR, YA, LT, GJM, GAJ and JAG designed research; RR, YA, LT, W-HC and JJAB performed research; RR, YA, LT and GAJ analyzed data; RR, YA, LT, GAJ and JAG wrote the paper.

## Acknowledgments

We would like to thank Dr. Yaniv Ziv for his expert guidance and advice. Eng. Anatoly Shapochnikov offered excellent technical support.

## Multimedia legends

**Movie 1. Synchronous spatiotemporal wave-like patterns in striatal cholinergic neuropil of a freely moving mouse.** Microendoscopic imaging of the DLS of a freely-moving mouse expressing the calcium indicator GCaMP6s selectively in CINs. The size of the visualized area is approximately 600 µm by 900 µm, and the movie is presented in real time. Imaging reveals fluctuations in the fluorescence of cholinergic neuropil characterized by rapid bursts of activation that permeate the entire field of view and decay slowly. The movie includes three examples of activation events, in which the neuropil activation propagates from top left to bottom right exhibiting a wave-like spatiotemporal structure.

**Movie 2. The cholinergic neuropil calcium signal precedes the somatic signal.** Surface plot visualization of the microendoscopic calcium imaging signal in the DLS of a freely-moving mouse (as in Movie 1). Visualized area is a 40 µm by 60 µm patch consisting of a single CIN soma surrounded by cholinergic neuropil. Example of an activation event demonstrating that somatic fluctuations are superimposed upon the temporal fluctuations of surrounding pixels and are slightly preceded by them.

**Movie 3. Wave-like propagation of activation in space.** Time reversed Δ*F*/*F* after appropriate scaling and shift (see Figure 3B, Materials and Methods). Various colors correspond to different time points in the propagation of the wave. Gray area marks the FoV. Movie is slowed down by a factor of 10.

**Movie 4. Analysis of synchronous wave-like propagation of cholinergic neuropil activation along dorsal surface of striatum for another mouse.** Left: Same as Movie 1. Neuropil activation propagates from bottom to top. Right: Map of time delays of the neuropil signal peaks (in each pixel) relative to the time of peak of the average annular signal, as in Figure 2D.

